# Computational and Structure-Guided E-Pharmacophore-Based Virtual Screening for the Identification of Novel NEK2 Kinase Inhibitors as Potential Anticancer Agents

**DOI:** 10.64898/2026.07.31.742111

**Authors:** Hafiz Muzzammel Rehman, Ayesha Latif, Hafiz Muhammad Hammad, Muhammad Sajjad

## Abstract

Cancer is a serious public health problem, and is becoming more common, with a projected increase in deaths and more than 25 million new cases in coming decades. A number of molecular mechanisms are involved in the tumoral process, with one of them, never in mitosis A–related kinase 2 (NEK2), a serine/threonine protein kinase, being a frequent target of amplification in various malignancies that is responsible for chromosomal instability, aneuploidy and activation of several oncogenic pathways. Available kinase inhibitors are not yet optimized in terms of their pharmacokinetic properties for clinical use, and current therapies, such as chemotherapeutic agents or immunotherapies are often limited by their resistance. In silico methods represent an effective tool to search for novel potent inhibitors, before testing in animals, with time constraints and limited resources. To find new inhibitors of NEK2, we used E pharmacophore–based modeling and structure based virtual screening in this study. NEK2 was chosen as the target for therapeutic intervention and an energy optimized pharmacophore model was employed to screen the Enamine REAL library of millions of compounds. Pharmacodynamic and Pharmacokinetic properties of the Top hits were tested using ADMET profiling. These were further screened using molecular docking (standard precision and extra precision) and virtual screening to obtain three lead compounds 1, 2, and 3 which have docking score of −7.414, −8.037 and −7.562 respectively. MM-GBSA calculations were used to estimate the binding free energies for these complexes, which were determined to be −54.92, −54.18 and −49.23 kcal/mol. Lastly, 100 ns molecular dynamics simulations have been run to evaluate complex stability in dynamic situations. The overall results of the MD showed the overall stability of the NEK2–ligand complexes, and thus these three compounds are promising NEK2 inhibitor candidates and could be further validated in vitro and in vivo for clinical application.

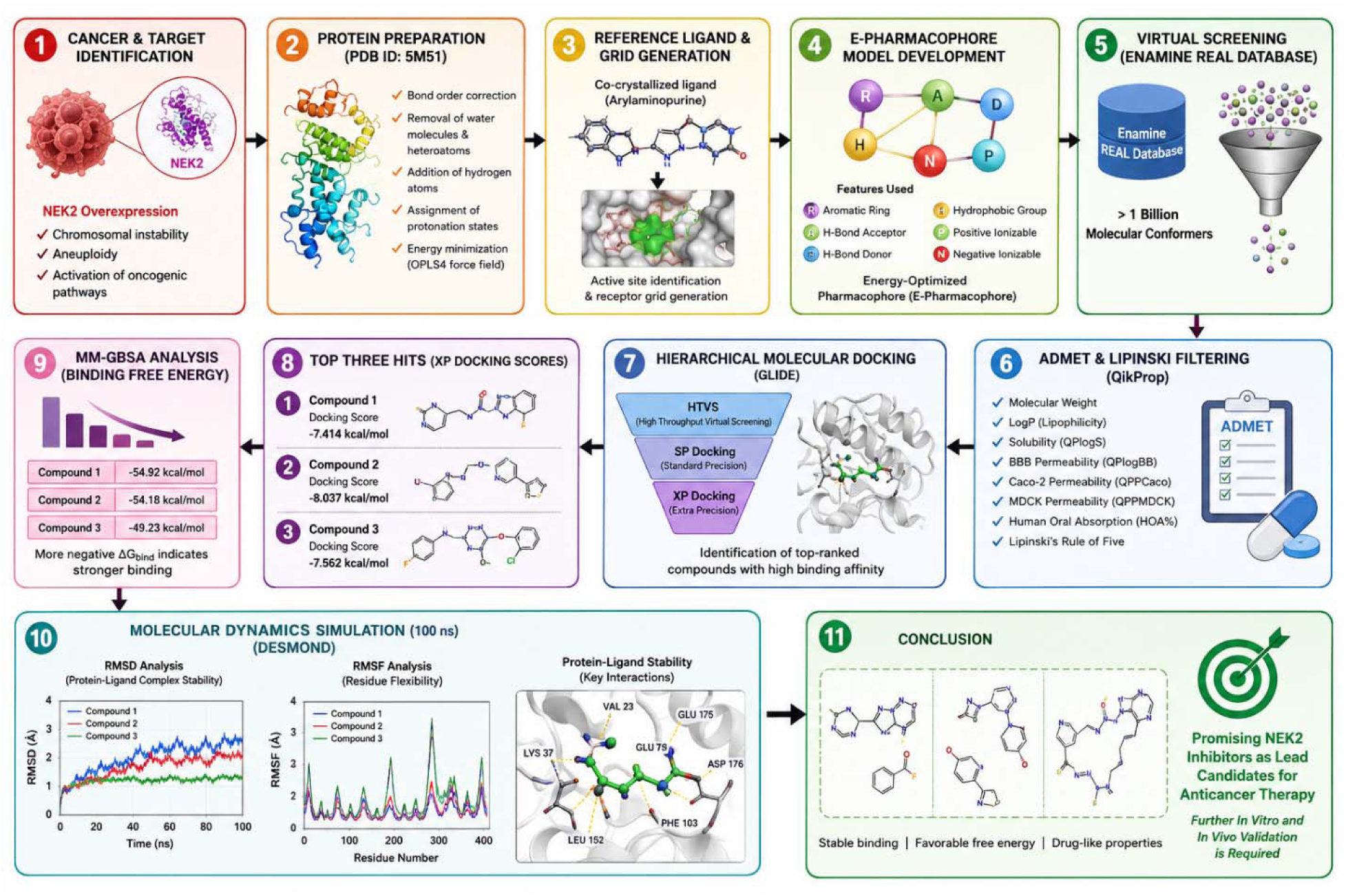

## 1. Introduction

Cancer is an important public health problem around the world, with approximately 20 million new cases and 9.7 million deaths in 2022 [1–4] and an estimated of up to 35 million new cases by 2050. Breast and lung cancers alone cause millions of cases and deaths and incidence is increasing most rapidly in transitioning, lower resource countries [5,6,1,7]. This growing burden is driven not only by demographics, but also by molecular drivers that promote aggressive disease and treatment resistance.

One such driver is Never in mitosis A-related kinase 2 (NEK2), a serine/threonine kinase essential for the separation of centrosomes and the segregation and progression of the mitoses [8]. NEK2 is always found to be overexpressed at protein and mRNA levels in a wide variety of tumors such as liver, breast, gastric, colorectal, esophageal, cervical and glioma cancers, as well as multiple myeloma [8–16]. By performing pan cancer analysis of 33 tumor types, NEK2 was found to be upregulated in nearly all cancers, with high expression levels significantly influencing overall, disease specific and progression free survival in many tumor entities [10]. In digestive system cancers and solid tumors, there are meta-analyses with a total of up to 4,897 patients that showed ∼1.5–2fold higher risk of death and recurrence in patients with NEK2 overexpression, including in hepatocellular, colorectal, lung, breast and glioma [15,16].

On the biological level, NEK2 overexpression causes chromosomal instability and aneuploidy and stimulates a number of oncogenic pathways such as the PI3K/AKT pathway, the Wnt/β catenin pathway, the MAPK pathway, and the NF κB pathway, which promote proliferation, invasion, metastasis and stem-like phenotypes, respectively [8,9,13,14,15]. It also has an impact on the tumor microenvironment: high NEK2 in gastric cancer associates with low immune cell infiltration and low cancer immunity cycle [11]; in multiple myeloma and pan cancer datasets NEK2 correlates with immune checkpoints, TMB and MSI, suggesting it may be an immune related biomarker and a target for immunotherapy [8,10] NEK2 upregulation has been associated with therapy resistance from a clinical perspective, for example in cervical cancer, whereby upregulation of NEK2 is correlated to resistance to cisplatin, and the induction of resistance to cisplatin in vitro and in vivo is associated with activation of the Wnt/β catenin pathway [9]; similarly, in glioblastoma and esophageal adenocarcinoma, there is a link between upregulation of NEK2 and growth, and genetic or pharmacological inhibition of NEK2 blocks tumor progression in vitro and in vivo [12,17,18,19].

Cancer treatments in current use include chemotherapy agents (such as the taxanes and platinum agents), targeted agents (such as EGFR inhibitors), immunotherapies (such as PD-1/PD-L1 inhibitors), and proteasome inhibitors (such as bortezomib) [20,21]. These strategies are, however, often impaired by the development of drug resistance (often related to the overexpression of NEK2) and by off-target toxicities. The inhibitors of NEK2 that are available demonstrate interesting preclinical results, but are not yet fully optimized for their pharmacokinetic and pharmacodynamic properties and are not yet fully selective [8]. Given the scale of the worldwide cancer burden and the pan-cancer association of NEK2 with poor outcomes, drug resistance and immune escape, there is a critical need for rationally designed, potent and selective NEK2 inhibitors that can be integrated into modern combination regimens.

A solution to this gap can be provided efficiently using structure based and ligand based computational approaches. The high-resolution NEK2 structures along with the increasing number of active scaffolds in the data base can be used in bioinformatics and chemo-informatics pipelines to create inhibitors with increased affinity and specificity and better drug like properties. The integrated in silico workflow including E pharmacophore modelling, virtual screening, molecular docking, ADMET prediction, MM GBSA binding free energy calculation and molecular dynamics simulations allows identification and prioritization of candidates at a high throughput level, without using time-consuming experimental methods [22,23]. This is particularly appropriate of a target such as NEK2 which has multi cancer, global clinical impact, but for which clinically viable inhibitors are lacking.

The aim of the present study is, therefore, to (i) systematically exploit the structural and pharmacophore information of NEK2 to generate an E pharmacophore model, (ii) perform large scale virtual screening to look for novel NEK2 directed chemotypes, (iii) characterize the binding modes and affinities using docking and MM GBSA, (iv) filter hits using in silico ADMET screening, and (v) evaluate the stability of top NEK2–ligand complexes by molecular dynamics simulations. The study aims to provide NEK2 inhibitors that will be applicable globally and potentially impact outcomes in a variety of high burden cancers.

## 2. Methodology

### 2.1. Protein Preparation

The Protein Data Bank (PDB), [24] provided the crystal structure of the Nek2 kinase in complex with an aryl aminopurine at 1.90 Å resolution (PDB ID: 5M51). The protein structure was first prepared and optimized via the Schrödinger suite of programs, Protein Preparation Wizard [25] before computational analysis. The procedure was carried out to remove structural inconsistencies and quality of the protein model was improved for further molecular docking and molecular dynamics studies.

The preparation of the proteins included the correction of bond orders, the removal of crystallographic water molecules and other irrelevant heteroatoms and the addition of hydrogen atoms. Moreover, appropriate tautomeric and ionization states of the amino acid residues were assigned to get a chemically correct structure [26] (Schrödinger Release 2018-1). To further preprocess the complex of the Nek2-inhibitor, water molecules within 5 Å of the hetero groups were removed. The missing hydrogen atoms were then added and the hydrogen-bonding network was optimized. Finally, the stable and energetically favorable protein structure was produced by performing restrained energy minimization with the OPLS4 force field for further analysis [27].

### 2.2. Ligand Preparation

The LigPrep module, as part of the Schrödinger software package, was used for ligand preparation. The module was used to create a new refined and customized library of ligands using suitable preparation filters. In the process, geometry optimization of the ligands was performed by optimization of the torsion angles and then their protonation states were assigned [32].

In addition, further refinements such as the insertion of hydrogen atoms, the generation of possible stereoisomeric forms, and the determination of relevant ionization states were made. All of the ligands prepared were then energy minimized using the OPLS 4 force field, built into the PHASE module [33] to find energetically optimized three-dimensional conformations. The following procedure allowed to generate low energy ligand structures suitable for further docking studies.

### 2.3. Molecular Docking

The interaction of the prepared ligands with the Nek2 protein was done with GLIDE module of the Schrödinger suite [28]. The molecular docking results indicated that several binding orientations of each molecule were obtained within the active site of Nek2, which allowed to assess their binding orientations and affinities. The docking results were further investigated employing more detailed information about the non-bonded interactions that underlie protein–ligand recognition, called Glide Extra Precision (XP) descriptors [29].

The binding pocket was assessed for the most favorable ligand conformations using pose scoring parameters from the Glide XP program, which measures the energetic contribution for each pose. The co-crystallized ligand (arylaminopurine) from the Nek2 crystal structure (PDB ID: 5M51) was used as the reference molecule to create a receptor grid with the Glide Grid Generation tool before the docking. This step helped to identify a docking box, which included the region of the active site of Nek2 [30].

After generating the grid, the simulation of the docking of the prepared ligand structures was carried out. The energy optimized pharmacophore (E-pharmacophore) model was then developed for the resulting protein–ligand complex that showed good binding.

### 2.4. The Energy-Optimized Pharmacophore (E-Pharmacophore) Modeling technique

To include the intrinsic nature of the target binding pocket a pharmacophore model was created but with energy optimization, E-pharmacophore. This method allows for energetically favorable hypotheses of pharmacophores to be developed and then can be used for the quick computational screening of millions of compounds. The E-pharmacophore methodology combines the idea of ligand-based pharmacophore with the structure-based information, which allows to identify the most important interactions between the ligand and the receptor, as well as to take into account the flexibility of the receptor active site [31].

Using the selected Nek2–ligand complex from the molecular docking studies, the E-pharmacophore model was created in the PHASE module in the Schrödinger suite. A set of six default features were selected for model generation: aromatic ring (R), negatively ionizable group (N), hydrogen bond acceptor (A), hydrogen bond donor (D), hydrophobic group (H) and positively ionizable group (P). These features were employed to develop an E-pharmacophore hypothesis that can be used to identify compounds that have favorable interaction patterns towards the Nek2 active site.

### 2.5. E-Pharmacophore-Based Virtual Screening

The E-pharmacophore model that was obtained from the Nek2–ligand complex was used as a query to perform virtual screening using the PHASE module of the Schrödinger suite. The screening was performed over Enamine REAL Database (RDB), which contains over 1 billion molecular conformers. This is a high-throughput virtual screening that enabled the swift discovery of chemicals with the pharmacophoric profile that complements Nek2’s binding site.

### 2.6. ADMET Evaluation and Lipinski Filtering

Using the QikProp module of the Schrödinger platform, the pharmacokinetic and drug-likeness properties of the compounds retrieved from the virtual screening based on the E-pharmacophore were assessed. The molecules that were screened underwent ADMET analysis to determine their potential as drug candidates and further screening based on Lipinski’s rule of five [34].

Molecular weight, blood–brain barrier partition coefficient, lipophilicity (log Po/w), parameters related to hydrophobicity and hydrophilicity (QPlogPo/w, QPlogS, QPPCaco, QPlogBB and QPPMDCK) were selected and examined. In addition, the percentage of human oral absorption (HOA%) was determined to evaluate the oral bioavailability potential of the selected compounds.

### 2.7. Virtual Screening based on molecular docking

After E-pharmacophore screening of the compounds in the Enamine REAL Database and filtering for ADME/T properties, the most promising compounds were short-listed for additional evaluation using molecular docking with Nek2. To systematically prioritize potential inhibitors, a hierarchical docking workflow was set up with the use of the GLIDE (Grid-Based Ligand Docking with Energetics) module of the Schrödinger suite.

All the compounds selected were then subjected to High Throughput Virtual Screening (HTVS) to quickly identify compounds that have good binding potential. The HTVS stage yielded the compounds that passed this filter that were further processed for Standard Precision (SP) protocol to get a better approximation of the binding interactions. Lastly, the top ranked molecules from the docking results with the SP were subjected to further refinement using the Extra Precision (XP) docking. This multi-stage screening system allowed the identification of top scoring compounds with good binding affinity to the Nek2 active site.

### 2.8. Estimating Binding Free Energy with MM-GBSA

Molecular Mechanics–Generalized Born Surface Area (MM-GBSA) calculations were then run to further investigate the binding strength of the top-ranked compounds identified by XP docking. This method gives a quantitative measure of the binding affinity of the target protein to the ligand molecules. The Prime module, included in the Schrödinger platform, was used to obtain a theoretical binding free energy calculation for the selected protein-ligand complexes which was derived from the docking results and the Glide scoring function [35] using the Schrödinger platform.

The MM-GBSA method estimates binding free energy based on the energetic contribution of the isolated protein, isolated ligand and the protein–ligand complex [36]. The free energy of binding ΔGbind was determined using the following equation:

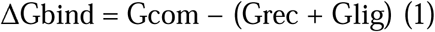

Here, Gcom is the free energy of a protein–ligand complex, Grec the free energy of the receptor, and Glig the free energy of the ligand.

The overall binding free energy consists of enthalpic and entropic terms [37] represented as:

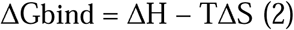

where ΔH is the enthalpic contribution, (T) is the absolute temperature and ΔS is the entropy change of the ligand binding.

### 2.9. Molecular Dynamics Simulations

To estimate the dynamic behavior, conformational stability and binding persistence of the selected protein-ligand complexes which were obtained from molecular docking studies, molecular dynamic (MD) simulations were performed in the Desmond module of Schrödinger suite [33]. All simulations started from the docked complexes.

The water models for each system were the Simple Point Charge (SPC) water model, and the overall charge neutrality was accomplished by the addition of appropriate Na+ and Cl− counterions. Before production simulations, the systems were energy minimized and relaxed under an NPT ensemble to eliminate unwanted steric contacts as well as excessive potential energy in the starting structures.

Next, MD simulations were carried out for 100 ns at a temperature of 300 K, and an NPT simulation at 1 atm with OPLS4 force field. Root mean square deviation (RMSD) was used to determine the stability of the protein–ligand complexes over the course of the simulation, while root mean square fluctuation (RMSF) was used to examine residue-level and structural fluctuations.

## 3. Results

### 3.1. Energy-Optimized Pharmacophore (E-Pharmacophore)

By generating the energy-optimized pharmacophore hypothesis, the key molecular characteristics that are crucial for binding into the active site of NEK2 kinase were identified. The pharmacophore model consisted of one hydrogen bond acceptor (A), two aromatic ring features (R) and three hydrophobic features (D) (Figure 1). The features are the important electrostatic and hydrophobic contacts that are involved in the recognition and stabilization of ligands in the ATP binding pocket of NEK2.

**Figure 1.**
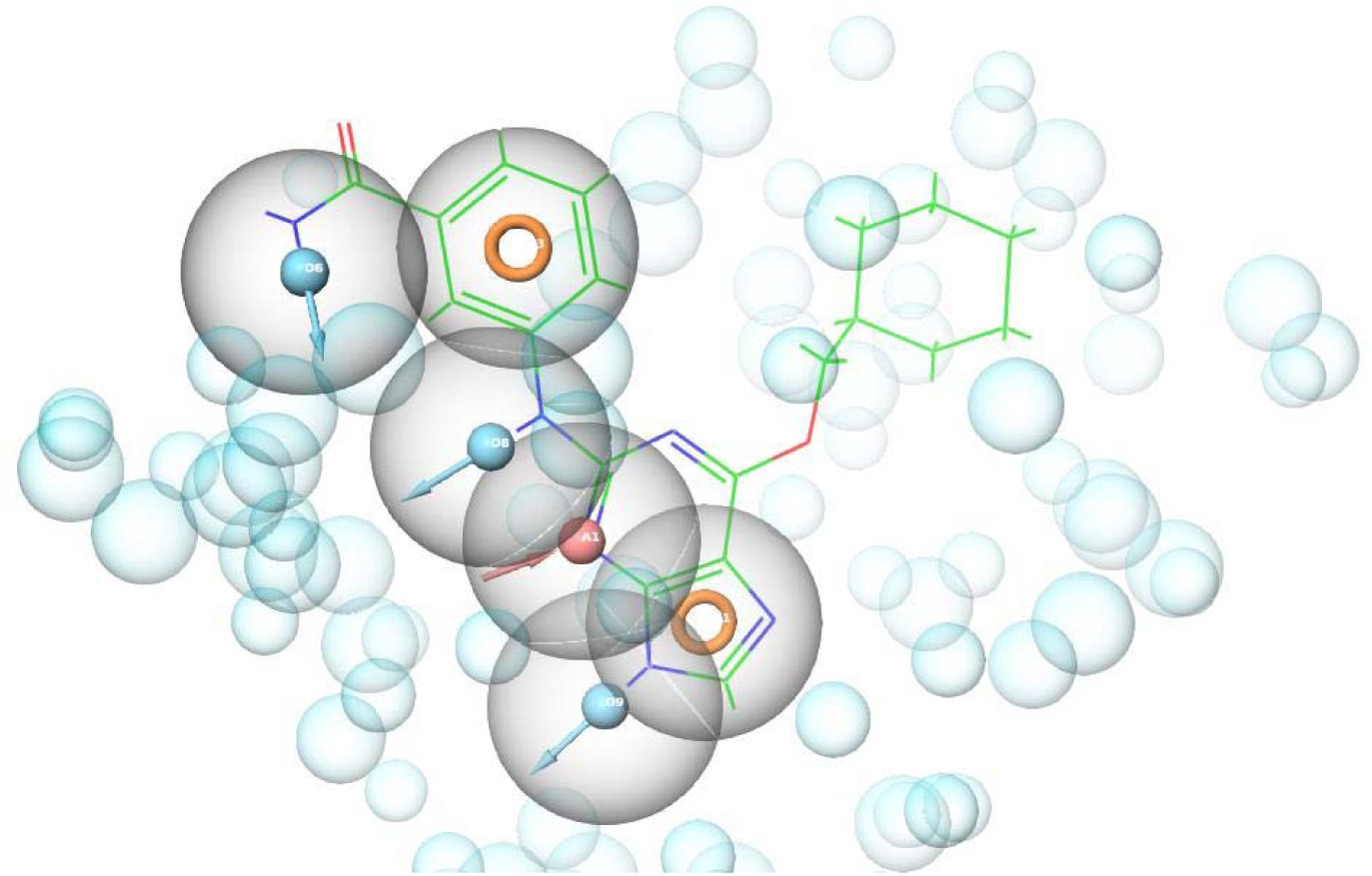
3D E-pharmacophore based model of NEK2, with one hydrogen bond acceptor (A1), two aromatic ring (R) features, and three hydrophobic (D6, D8 and D9) features. The excluded volumes for *steric restraints shown by the light blue spheres correspond to the position of the reference ligand (green structure) in the generation of the pharmacophore*

The hydrogen bond acceptor feature shows the role of hydrogen bonding with active-site residues, and the aromatic ring feature shows the role of aromatic interaction in ligand binding. In addition, the hydrophobic characteristics indicate potential nonpolar contacts in the kinase binding cavity, which will be essential for improving binding affinity. The pharmacophore model generated also included excluded volume spheres that specify the steric limitations of the binding pocket of NEK2 and which help to sort compounds by space.

The 3D pharmacophore hypothesis was then used in virtual screening to find compounds with the desired structural and physicochemical properties to be used as potential inhibitors of NEK2.

### 3.2. ADMET Analysis and Lipinski Parameters

Selected lead compounds pharmacokinetics profile was assessed using ADME/T descriptors and the results shown in Table 1. All three compounds had good drug-like properties with molecular weight range from 322.366 to 432.524 Da and QPlogPo/w 2.800 to 3.017, which is considered as balanced lipophilicity. Proposed human oral absorption was between 78.506% to 100% indicating good oral bioavailability.

**Table 1.**
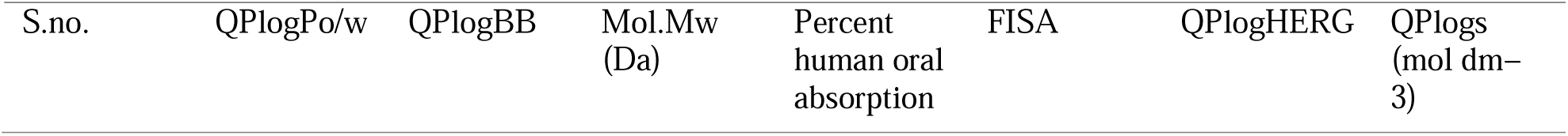
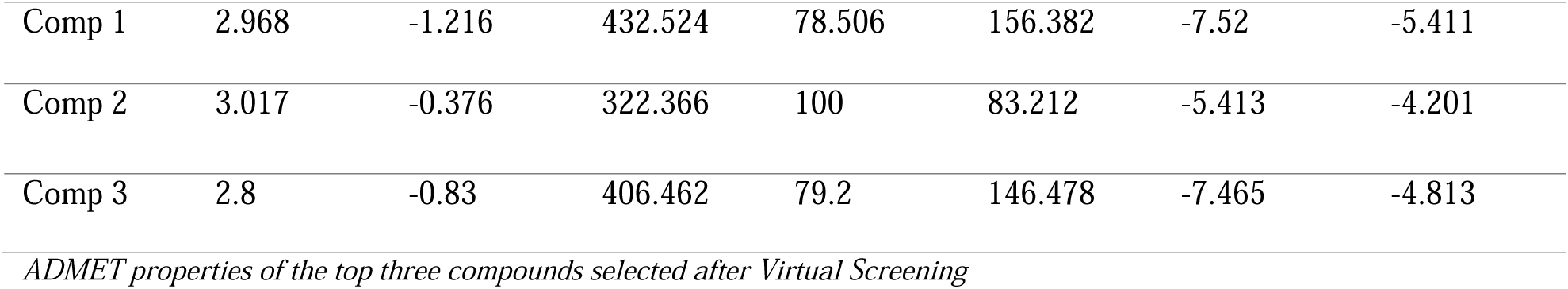

The values of QPlogS were acceptable for aqueous solubility and QPlogBB were indicative of limited penetration of the blood-brain barrier. Moreover, the QPlogHERG values were also in safe parameter ranges, suggesting low potential for cardiotoxicity. Among the three top-hits, Compound 2 had the best pharmacokinetic profile, with a predicted oral absorption of 100% and a low molecular weight. In conclusion, the ADMET analysis validated that all the selected compounds have good drug-like properties and can be explored as NEK2 inhibitors.

### 3.3. Structure-Based High Throughput Virtual Screening using Receptor Models

Pharmacophore-based screening and ADMET filtering were performed to retrieve the compounds which were subjected to receptor-based virtual screening against the active site of the NEK2 receptor. The compounds with highest docking scores, drug-likeness properties and binding interactions were chosen as top-ranked compounds. Three lead compounds were found to show good binding affinity towards NEK2 with XP Glide scores ranging from −7.577 to −8.392 kcal/mol (Table 2).

**Table 2:** Compound codes, XPG Scores, Docking Scores, Rule of five violations and other stability parameters obtained through Molecular docking and MM-GBSA analysis. Top hits are arranged in the descending order of MM-GBSA.

| Compounds | Compound Code | XPG Score | Mol MW | MM-GBSA | Docking Score | Rule of five violation | Prime Energy |
| --- | --- | --- | --- | --- | --- | --- | --- |
| Comp 1 | 2531 | -8.392 | 394.519 | -54.92 | -7.414 | 0 | -10920.2 |
| Comp 2 | 936 | -8.038 | 322.366 | -54.18 | -8.037 | 0 | -10955.5 |
| Comp 3 | 3010 | -7.577 | 406.462 | -49.23 | -7.562 | 0 | -10910 |

Of the hits the highest binding affinity was exhibited by Compound 1 (XP Glide score of −8.392 kcal/mol) and docking score of −7.414kcal/mol). Compound 2 also showed good binding affinity, XP Glide: (−8.038 kcal/mol) and docking score: −8.037 kcal/mol. The XP Glide score was calculated to be −7.577 kcal/mol for compound 3, which is a relatively low but still good binding affinity. The selected compounds met the Lipinski’s rule of 5 with no violation, and showed good PK properties, which are considered as good indicators for potential as NEK2 inhibitors. Hence, these lead compounds (Table 3) were chosen and subjected to further MM-GBSA and molecular dynamics simulation studies.

**Table 3:**
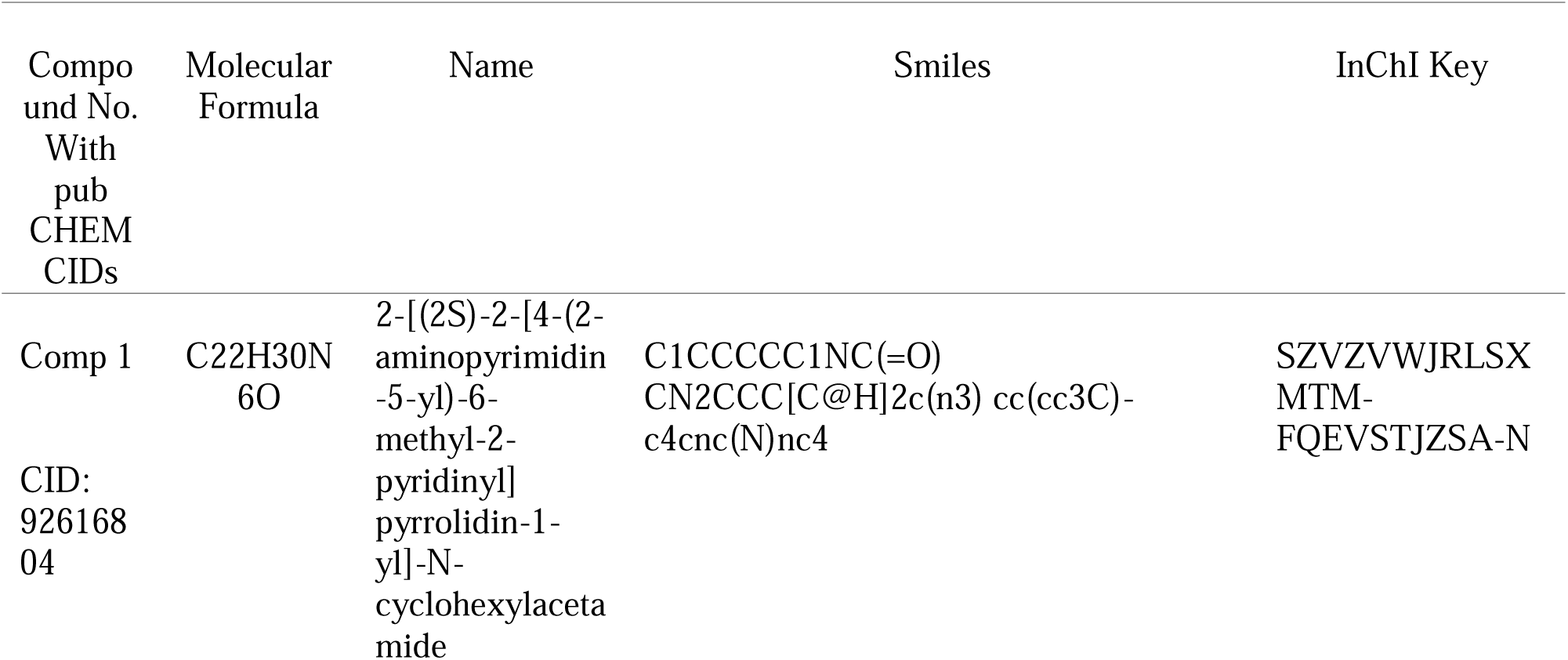
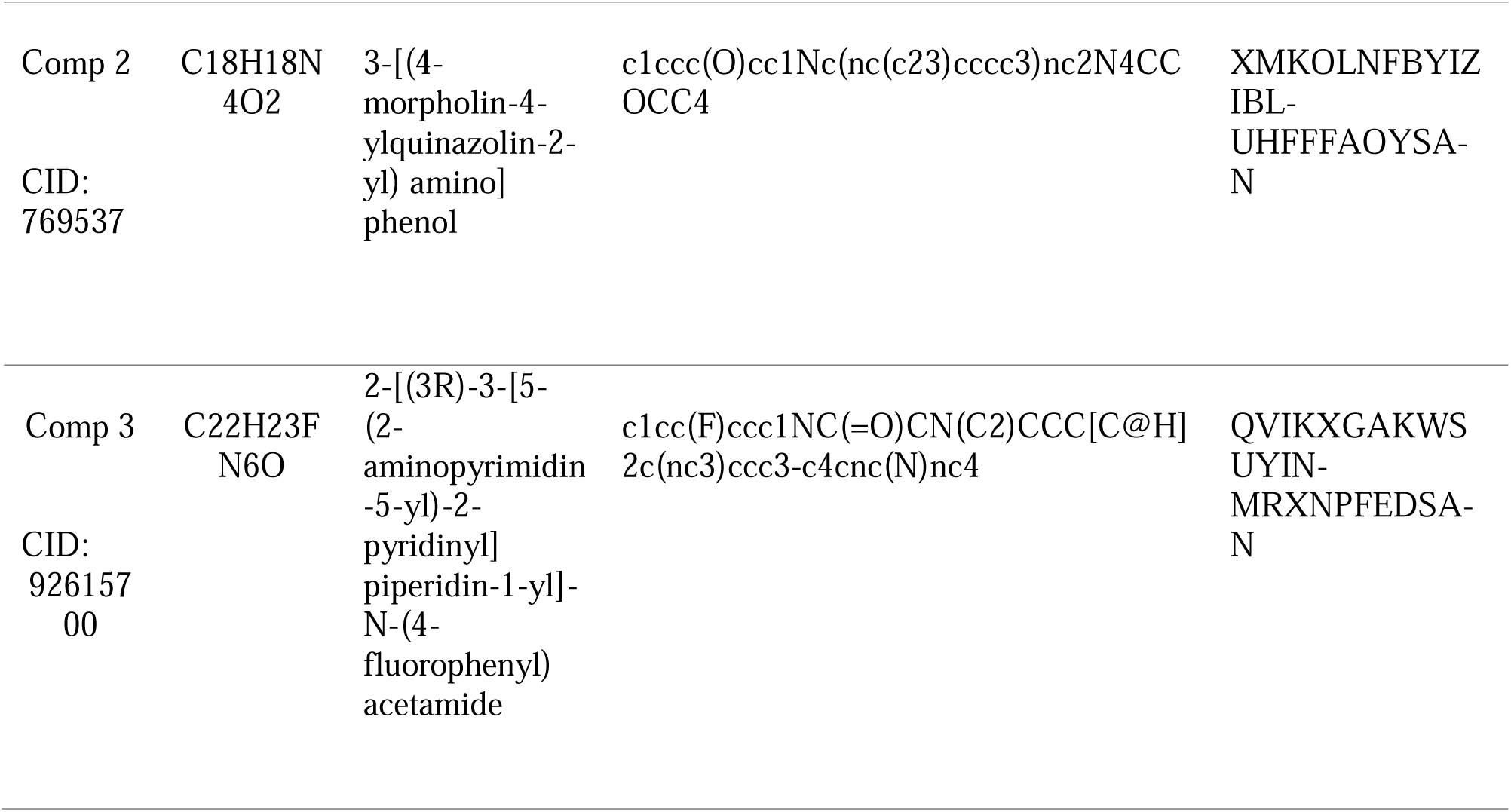
The molecular formula, PubChem Compound IDs, IUPAC names, Smiles and InChI Key of the top three compounds.

### 3.4. Estimation of Binding Free Energies Using MM-GBSA

The binding free energies of the selected lead compounds were evaluated using MM-GBSA analysis to further assess their binding stability within the NEK2 active site. The molecular docking results were also validated by the binding free energies obtained for all three compounds (Table 2), which were found to be favorable.

The best binding free energy value was obtained for Compound 1 (−54.92 kcal/mol) followed closely by Compound 2 (−54.18 kcal/mol). In addition, compound 3 showed good binding stability with the MM-GBSA score of −49.23 kcal/mol. These compounds have favorable MM-GBSA values in addition to high XP Glide scores, suggesting their stable binding to the NEK2 binding pocket. Compounds 1–3 were identified as good lead candidates based on these results, and were selected for further molecular dynamics simulation studies.

### 3.5. Interaction profiling of Hit Compounds with NEK2 Binding Site Residues

The interaction pattern of the selected hit compounds with NEK2 was analyzed with the Schrödinger ligand interaction diagram, which reveals important intermolecular contacts such as hydrogen bonds and π–π stacking interactions, salt bridges, and other non-covalent interactions. [38] Of these, hydrogen bonds play especially critical roles in maintaining stability and specificity of protein–ligand complexes. [39]

In the case of NEK2 receptor, Compound 1 formed a hydrogen bond and a salt bridge with ASP_93 at distances of 1.94 Å and 4.10 Å, respectively. Further, a π–π stacking interaction was found with PHE_148 at a distance of 4.93 Å. There were also hydrogen-bond interactions between the ligand and GLY_87 at 2.13 Å and MET_86 at 2.11 Å (Figure 2(a))

**Figure 2.**
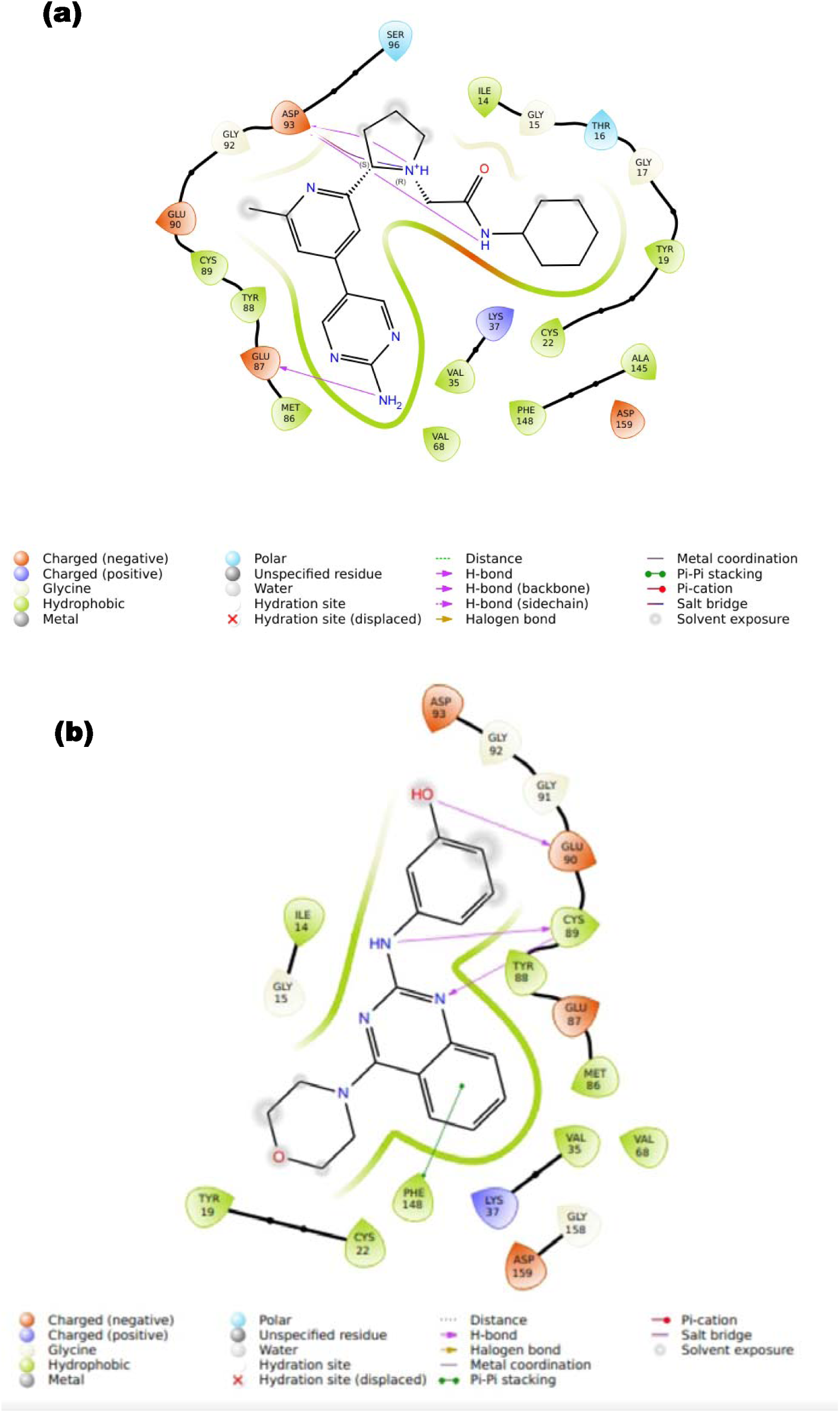
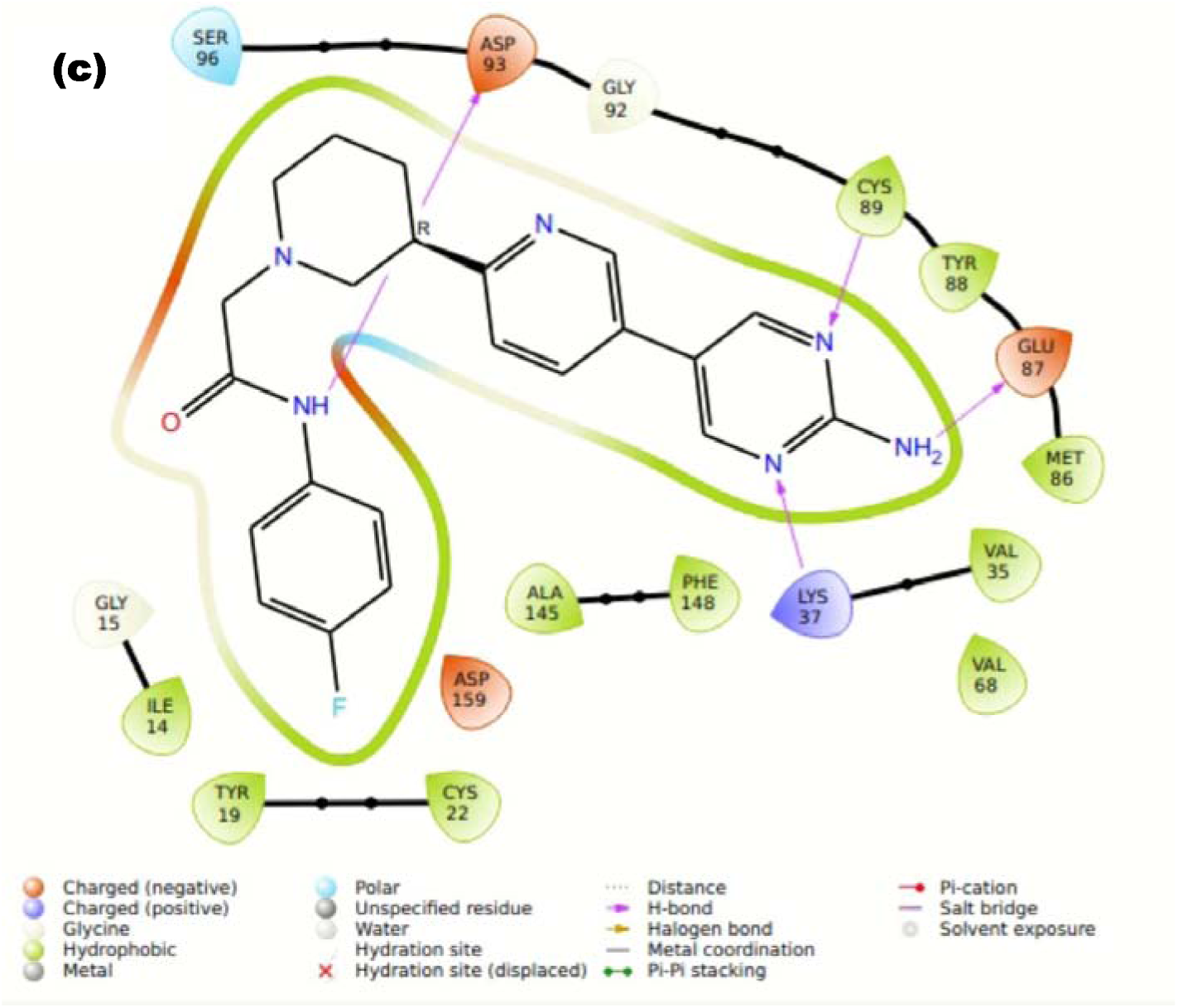
2D interaction maps of the top three hit compounds with NEK2 serine/threonine protein kinase: (a) intermolecular interactions between compound 1 and NEK2 binding residues; (b) intermolecular interactions between compound 2 and NEK2 binding residues; (c) intermolecular interactions between compound 3 and NEK2 binding residues.

The phenolic hydroxyl, linker NH and heterocyclic N atoms in compound 2 formed three hydrogen bonds with Glu90, Cys89 and Tyr88, respectively. It was found that there was a π–π stacking interaction with Phe148. In addition, hydrophobic interactions were observed with Ile14, Tyr19, Cys22, Met86, Val35, Val68, Tyr88, Cys89 and Phe148; and hydrogen bonding interactions were seen with Lys37, Glu87, Glu90, Asp93 and Asp159 which form the surrounding charged environment of the binding pocket. (Figure 2(b))

Four hydrogen bonds were formed between compound 3 and Asp93, Cys89, Glu87 and Lys37, the piperidine–pyridine junction region, pyrimidine N atoms and the exocyclic amino group of the aminopyrimidine part of the molecule. In addition, Ile14, Tyr19, Cys22, Ala145, Phe148, Val35, Val68, Met86, Tyr88 and Cys89 formed the surrounding hydrophobic environment of the binding pocket, while Asp93, Glu87, Lys37, Ser96 and Asp159 formed the surrounding polar or charged environment of the binding pocket. (Figure 2(c))

### 3.6. Molecular dynamics simulations

The dynamic behaviors and stabilities of the NEK2–ligand complexes were analyzed by molecular dynamics (MD) simulations over time. Three top compounds, which were chosen based on MM-GBSA binding free-energy calculations, were simulated for 100 ns. These compounds were Compound 1, Compound 2, and Compound 3 with binding free energies of −54.90, −54.18, and −49.23 kcal/mol respectively.

Conformational stability of each NEK2–ligand complex was evaluated by the root mean square difference (RMSD) analysis, and the flexibility and mobility of the residues were analyzed by root mean square fluctuation (RMSF) analysis. The values of RMSF also stayed below 3 Å in all the systems, showing the stability of the protein structure throughout the simulation. Similarly, RMSD values were mostly less than 3 Å indicating minor conformational changes from the starting structures.

During the entire simulation, the RMSD for Compound 1 was within 3 Å. A small deviation of about 0.4 Å was seen around 18 ns, but the system quickly settled into a stable state and stayed that way for the rest of the trajectory with no significant deviations. (Figure 3(a)) Compound 2 also showed good stability over the course of the simulation. A slight rise in RMSD was observed from about 35 to 42 ns after which a stable trajectory was observed as the complex reached equilibrium again. The RMSD values of the protein backbone were kept between 2.4 Å and 2.6 Å and the RMSD values of the ligand were kept below 2.0 Å, suggesting high level of retention of the ligand in the binding area. (Figure 3(b)) In the case of Compound 3, the major variations were not seen in the simulation, indicating that the complex was very stable. The RMSD of the ligand did not change through the 1st 95ns of the trajectory. A small increase was observed after 95 ns, presumably due to small conformational changes, but the system rapidly re-equilibrated. The RMSD values for NEK2 backbone structure were found to be at an average of 1.5 Å to 2.2 Å during 100 ns simulation. The data taken together shows that Compound 3 does bind steadily in the active site of NEK2. (Figure 3(c))

**Figure 3.**
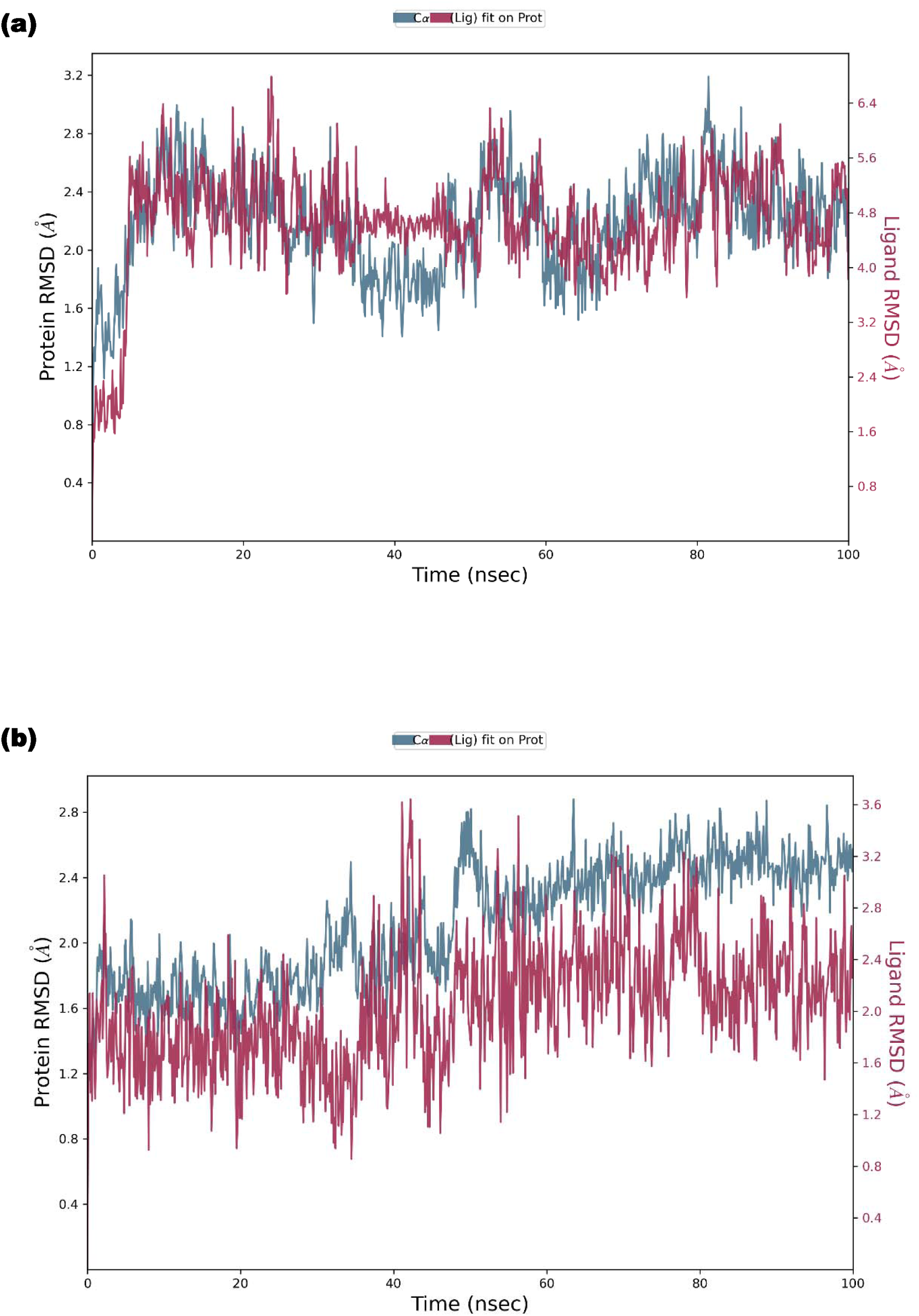
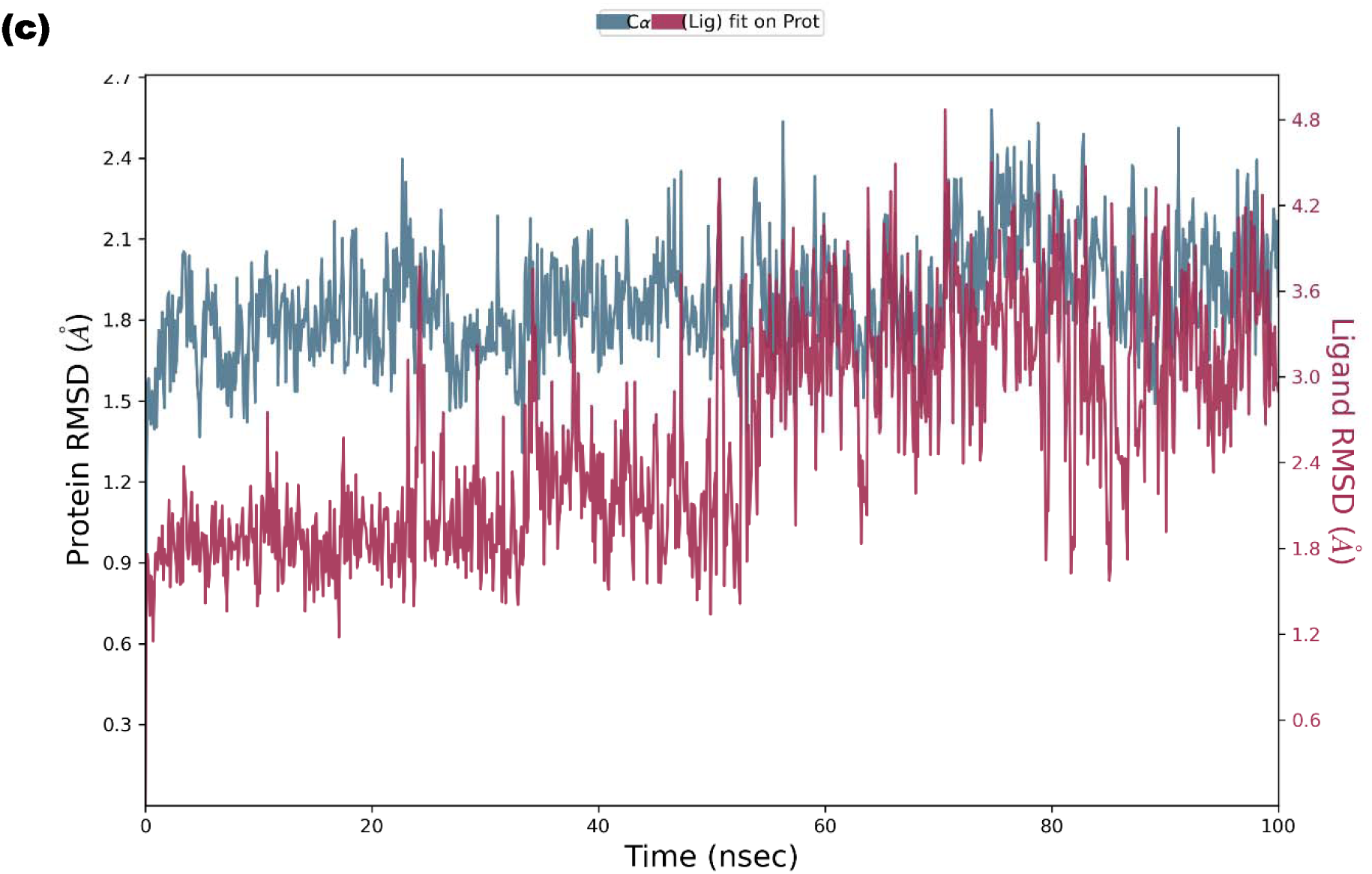
Root mean square deviation (RMSD) plots generated by the Desmond module of Schrödinger for 100ns: (a) RMSD plot of compound 1; (b) RMSD plot of compound 2; (c) RMSD plot of compound 3.

However, further analysis of the residue’s flexibility showed that the catalytic region of NEK2 in all three complexes is structurally stable. The root-mean-square fluctuations (RMSF) of RMSF profiles obtained from the simulated trajectories revealed no significant variations across the binding-site residues. In the RMSF plots, green vertical lines represent residues that participate in the binding of the ligand. (Figure 4 (a-c))

**Figure 4.**
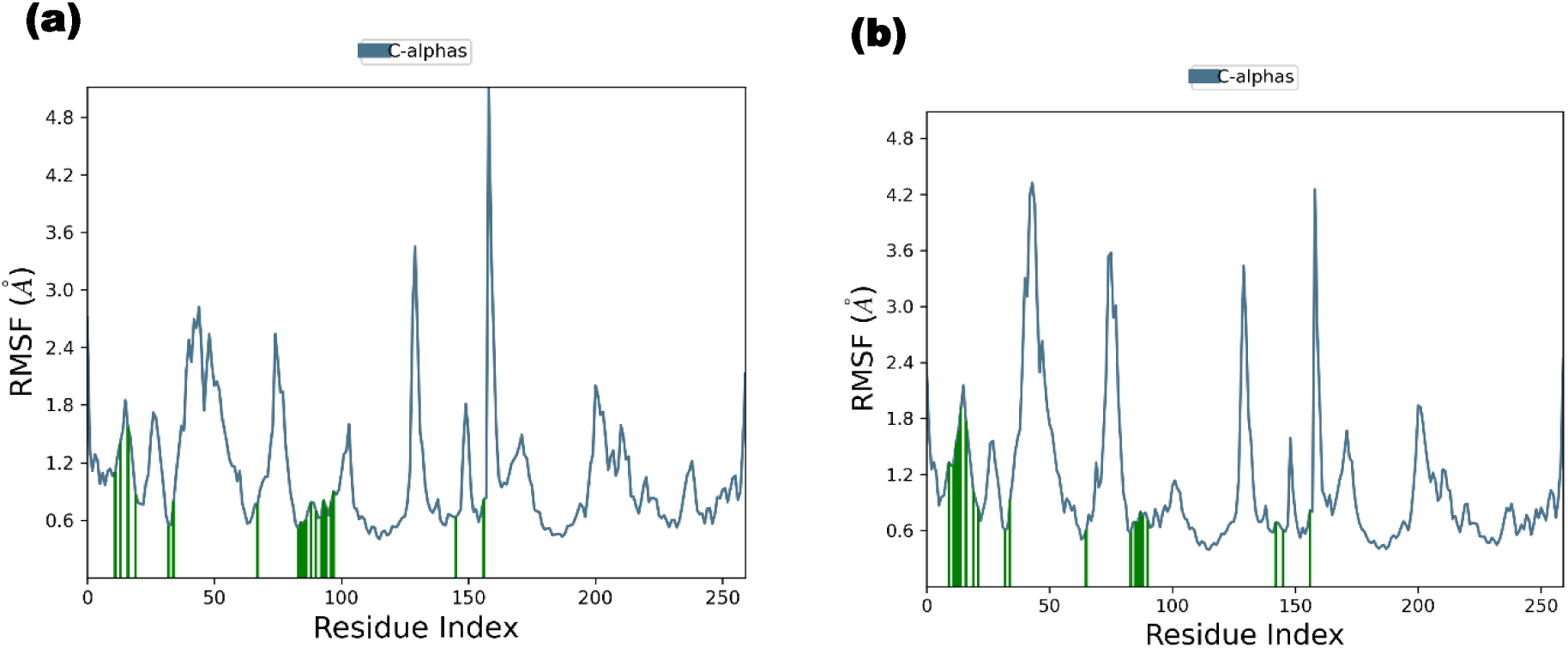
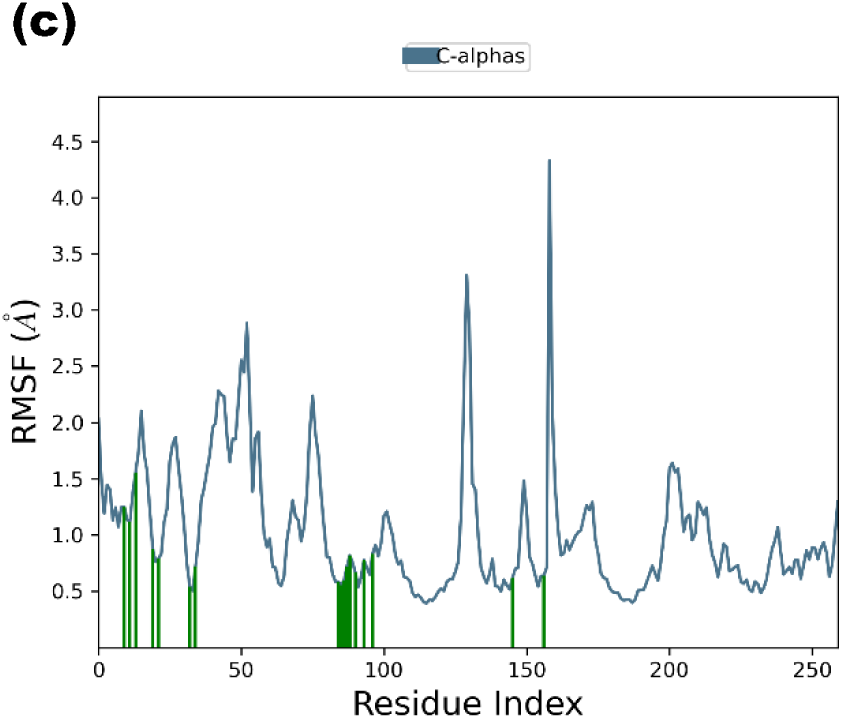
Root mean square fluctuations (RMSF) plots generated by the Desmond module of Schrödinger for 100 ns for (a) compound 1, (b) compound 2, and (c) compound 3 with NEK2 serine/ threonine protein kinase

In the MD simulation, the residues which interacted with the protein were studied and it was found that the interactions were stable for all three compounds with several important residues at the active site. In Compound 1, the most interacting residues were Glu87, Lys37, Tyr88, Cys89, Phe148, Asp93, Tyr19, Gly91, Asp159 and Met86. In Compound 2, the interacting residues included Tyr12, Phe148, Ile14, Val35, Lys37, Met86, Thr16, Cys22, Lys24, Cys89, Tyr88, Gly91, Asp93, Asp159 and Tyr19, whereas Compound 3 maintained stable contacts with Tyr12, Phe148, Val35, Cys22, Lys24, Thr99, Lys37, Tyr88, Cys89, Ile14, Thr16, Asp159 Gly91, and Asp93, throughout the simulation. Residues Tyr88, Cys89, Gly91, Asp93, Phe148 and Asp159 are recurrently involved in stabilizing the binding of the ligand in the active site of NEK2, indicating their crucial role in the binding of the ligand. (Figure 5 (a-c))

**Figure 5.**
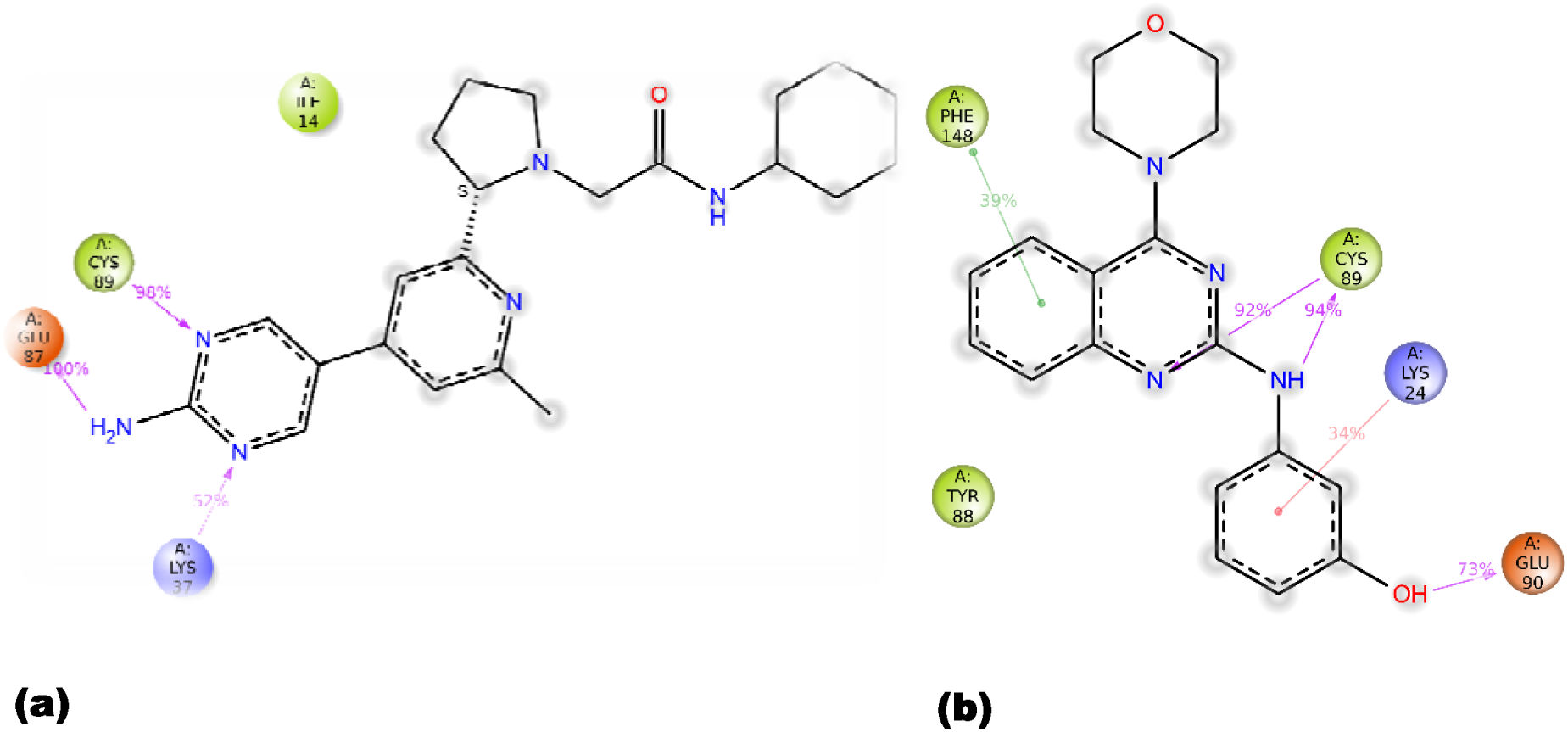
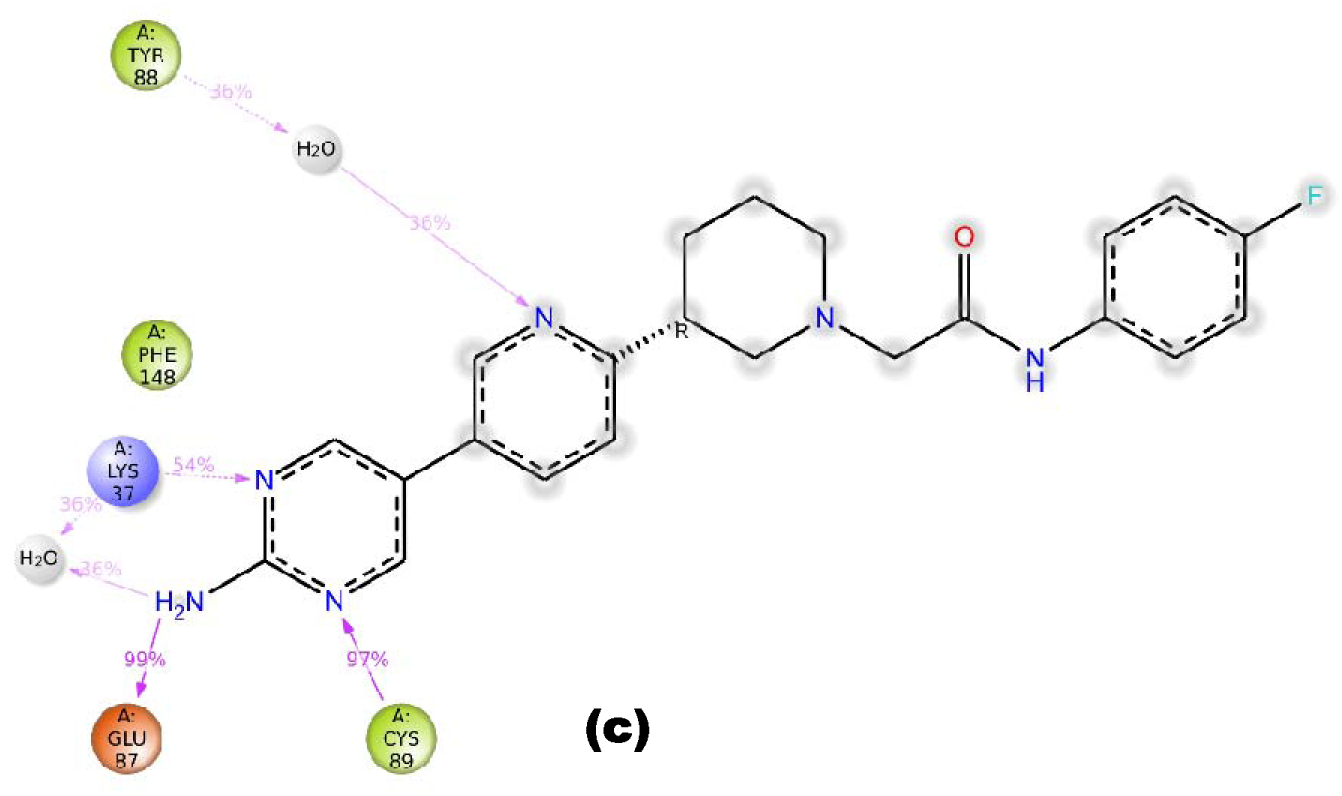
Schematic representation of detailed ligand atom interactions with protein (NEK2) residues that occur more than 30% of the simulation time in the selected trajectory (0-100ns) for (a) Compound 1, (b) Compound 2, and (c) Compound 3.

Overall, the MD simulation results showed that all three compounds maintained stable complexes with the NEK2 kinase (PDB ID: 5M51) throughout the simulation time (100 ns). The intermolecular interactions were also maintained during the simulation, along with minimal fluctuations in the structure and low RMSD and RMSF profiles, which all point to a strong binding affinity of the NEK2 active site. As such, such compounds can have the ability to successfully occupy and inhibit the active binding site of NEK2.

## 4. Discussion

In the present study, a virtual screening pipeline, which involved E-pharmacophore modeling, multistage docking, MM-GBSA, ADMET filtering, and 100 ns MD simulation was used to explore novel small-molecule inhibitors of NEK2. The three compounds with good XP docking score, negative MM-GBSA binding free energy, good drug-likeness characteristics and stable binding in the ATP pocket have the potential to be good lead compounds for the inhibition of NEK2.

Such integrated computational workflows have been shown to be effective in the previous drug discovery studies. To discover six potential SARS-CoV-2 RdRp inhibitors, Rehman et al used an E-pharmacophore-based virtual screening approach to the Enamine REAL database followed by HTVS virtual screening, molecular docking, MM-GBSA calculations, and MD simulations to determine the binding free energy and the stability of the protein–ligand complexes.[50] Likewise, Rehman et al. performed an E-pharmacophore-guided virtual screening of a large library of compounds against the NS2B–NS3 protease of the ZIKV and selected three compounds with good binding properties and pharmacokinetic parameters and with good dynamic stability as potential lead compounds.[51] The results of these studies show the reliability of the pharmacophore modeling, docking, free-energy prediction and MD simulation methodology for the discovery of biologically relevant lead compounds and validate the robustness of the approach used in the present study.

The docking results showed that all three ligands docked in the ATP-binding site, and the residues of the hinge region (MET86/GLY87/GLU87/CYS89), ASP93, PHE148, LYS37 and ASP159 (DFG-motif), were involved in key functional interactions with all three ligands. The contacts with the hinge region and the ASP93 and PHE148 ionic/π–π interactions are common in other NEK2 inhibitors that have been characterized, and these contacts have been found to be crucial for binding affinity and specificity. [38–40] This agreement is consistent with the structural plausibility of the predicted binding modes.

High docking scores were obtained with the MM GBSA binding free energies ranging from approximately –49 to –55 kcal/mol, which indicates good thermodynamics for the ligand binding. [41,42] MD simulations also showed that the complexes of NEK2 with inhibitors were dynamically stable; the RMSD of proteins and ligands were generally low (RMSD<3 Å) and RMSF values were low, suggesting that binding of the inhibitors did not lead to large destabilizing shifts. The stable kinase–inhibitor complexes from the virtual screening were further confirmed by similar RMSD/RMSF profiles with stable interactions with the catalytic and hinge region residues for 100 ns. [42,43]

The identified compounds were also found to have good ADMET properties and were also found to be drug-like and had good human oral absorption upon previous virtual screening studies of NEK2 and kinases [41,44] that filtered for drug likeness and human oral bioavailability, respectively, suggesting that our hits could be expected to be active in anticancer experimental tests.

These results are biologically significant, since NEK2 is found to be overexpressed in various tumors, associated with poor prognosis and associated with chromosomal instability, drug resistance and immune escape [39,45,46] and therefore, targeting NEK2 is a promising approach to improve the efficacy of chemotherapy, targeted therapy and immunotherapy. [46,47]

However, there are some drawbacks to this work. It is fully computational with no experimental data of IC□□, selectivity, or cellular activity data yet. Previous virtual screening studies about NEK2 showed that in silico studies require cell based and biochemical validation, kinase selectivity (versus PLK1), otherwise it can limit development. [40,41] Future studies need to be directed towards the synthesis of the three in silico hits, measure their inhibitory properties against the NEK2 kinase, evaluate their anticancer and chemo-sensitizing properties in the NEK2 overexpressing models, and optimize their PK and toxicity properties.

## 5. Conclusion

In this study, a multistage virtual screening approach, combined with molecular docking, MM-GBSA binding free-energy calculations, ADME/T analysis, and molecular dynamics simulation, were used to discover novel compounds with a target to NEK2. Three drug-like hit molecules were found that bind with good binding affinities, have stable interaction networks and good pharmacokinetic properties and remained stable for 100 ns. The identified compounds are potential candidate scaffolds that possess important structural and interaction characteristics of the NEK2 inhibitors. The role of NEK2 in tumor progression and therapeutic resistance makes these compounds attractive candidates to be used as starting points for designing NEK2 inhibitors for use in cancer therapy. However, their biological activity and therapeutic potential have to be validated in experimental studies such as enzymatic inhibition assays, cellular validation and selectivity profiling.

